# PatHoST: weakly supervised reconstruction of pathogen spatial distributions from host transcriptomes

**DOI:** 10.64898/2026.10.26.759336

**Authors:** Federico Di Marco, Francesca Nicola, Gaia Saldarini, Daniela Maria Cirillo, Nicola Ivan Lorè, Alex Graudenzi, Stefano de Pretis

## Abstract

Dual host–pathogen spatial transcriptomics enables spatially resolved investigation of infection, yet the completeness of pathogen detection remains poorly characterized. Here, we generated pathogen-only spatial transcriptomic sections from a chronic Mycobacterium abscessus lung infection model to experimentally benchmark pathogen detection, revealing substantially increased pathogen transcript recovery compared with conventional dual profiling. Building on these data, we developed PatHoST, a weakly supervised computational framework that uses pathogen-derived information to generate silver-standard labels and learns host transcriptional programs predictive of pathogen presence. PatHoST reconstructed pathogen spatial distributions across independent tissue sections and enabled the identification of localized host regulatory programs beyond conventional cluster-based analyses. Application to human SARS-CoV-2 lung spatial transcriptomics supported generalization across host species and pathogen classes. Together, PatHoST combines purpose-built experimental benchmarking with weakly supervised inference to recover latent pathogen information and extend the analytical capabilities of dual host–pathogen spatial transcriptomics.

## Introduction

Spatial transcriptomics (ST) has transformed the study of the tissue organization by enabling genome-wide gene expression profiling while preserving spatial context [1]. Recent advances in probe-based assays have further extended ST to infectious diseases by enabling the simultaneous detection of host and pathogen transcripts within the same tissue section [2, 3]. These dual host-pathogen spatial transcriptomics (dualST) approaches provide an unprecedented opportunity to investigate microbial niches together with the surrounding host response in situ. Despite these technological advances, pathogen transcripts are difficult to detect due to several technical limitations, such as poor permeabilization of the bacterial membrane or low RNA abundance and stability [4, 5], resulting in incomplete spatial maps that underestimate the true spatial distribution of infection. New experimental approaches are needed to overcome these issues with the aim to improve the characterization of host–pathogen interactions, pathogen transcriptional programs, and tissue-level infection dynamics.

Current computational approaches have primarily focused on improving transcript recovery through spatial smoothing, denoising, or latent representation learning [6–8]. While these methods effectively address technical noise in host transcriptomic data, they generally assume that the observed molecular signal provides an adequate representation of the underlying biology. In dualST datasets, however, the absence of detected pathogen transcripts cannot be interpreted as evidence of pathogen absence. Instead, technical dropouts create an inherently incomplete labeling problem, in which infected locations frequently appear indistinguishable from uninfected tissue.

This observation naturally reframes pathogen localization in ST as a weakly supervised (or positive–unlabeled) learning problem [9]. Rather than treating sparse pathogen counts as absolute ground truth, computational models can leverage the broader and more robust host transcriptional response, which provides an indirect yet biologically informative readout of pathogen activity. Because host responses extend beyond the relatively few locations where pathogen transcripts are directly detected, they offer a richer substrate for learning the spatial distribution of infection.

Here, we present PatHoST (Pathogen–Host Spatial Transcriptomics), a computational framework that reconstructs pathogen spatial distributions by learning the relationship between host transcriptional states and sparse pathogen observations. PatHoST first enhances pathogen signal through spatial projection and smoothing to generate silver-standard labels, and subsequently trains machine learning models using host gene expression alone to predict pathogen presence. Unlike conventional supervised approaches, PatHoST explicitly models the uncertainty associated with pathogen detection and leverages weak supervision to recover latent pathogen spatial distributions that remain only partially detected in the original data.

To establish a controlled benchmark for evaluating latent pathogen signal recovery, we generated dedicated spatial transcriptomic sections profiled exclusively with pathogen-specific probes. These pathogen-only acquisitions provides a higher-sensitivity experimental reference for pathogen detection, enabling the assessment of latent pathogen signal recovery beyond that achieved with conventional dualST assays. Using bacterial and viral infection datasets spanning different hosts and technologies, we show that PatHoST accurately reconstructs pathogen spatial distributions, generalizes across biological systems, and identifies host regulatory programs associated with infection that are not apparent using conventional analytical approaches.

By integrating host transcriptional responses, pathogen observations, and spatial information, PatHoST enables (i) improved pathogen detection and quantitative characterization by learning host response signatures, (ii) retrospective analysis of datasets with limited or absent pathogen capture, and (iii) a biological interpretation of host transcriptional programs associated with infections. Together, these results demonstrate that the host transcriptome can serve as informative proxy for pathogen spatial distribution, extending the utility of ST for investigating microbial virulence and host-pathogen interactions even when direct pathogen detection is incomplete.

## Results

### PatHoST overview

We developed PatHoST, a computational framework that leverages host gene expression to recover latent pathogen signal from dualST experiments. Starting from paired host and pathogen expression matrices, PatHoST integrates spatial signal enhancement, weakly supervised learning, and automated model selection to estimate the predicted likelihood of pathogen presence and reconstruct the corresponding predicted pathogen spatial distribution (Figure 1).

**Figure 1:**
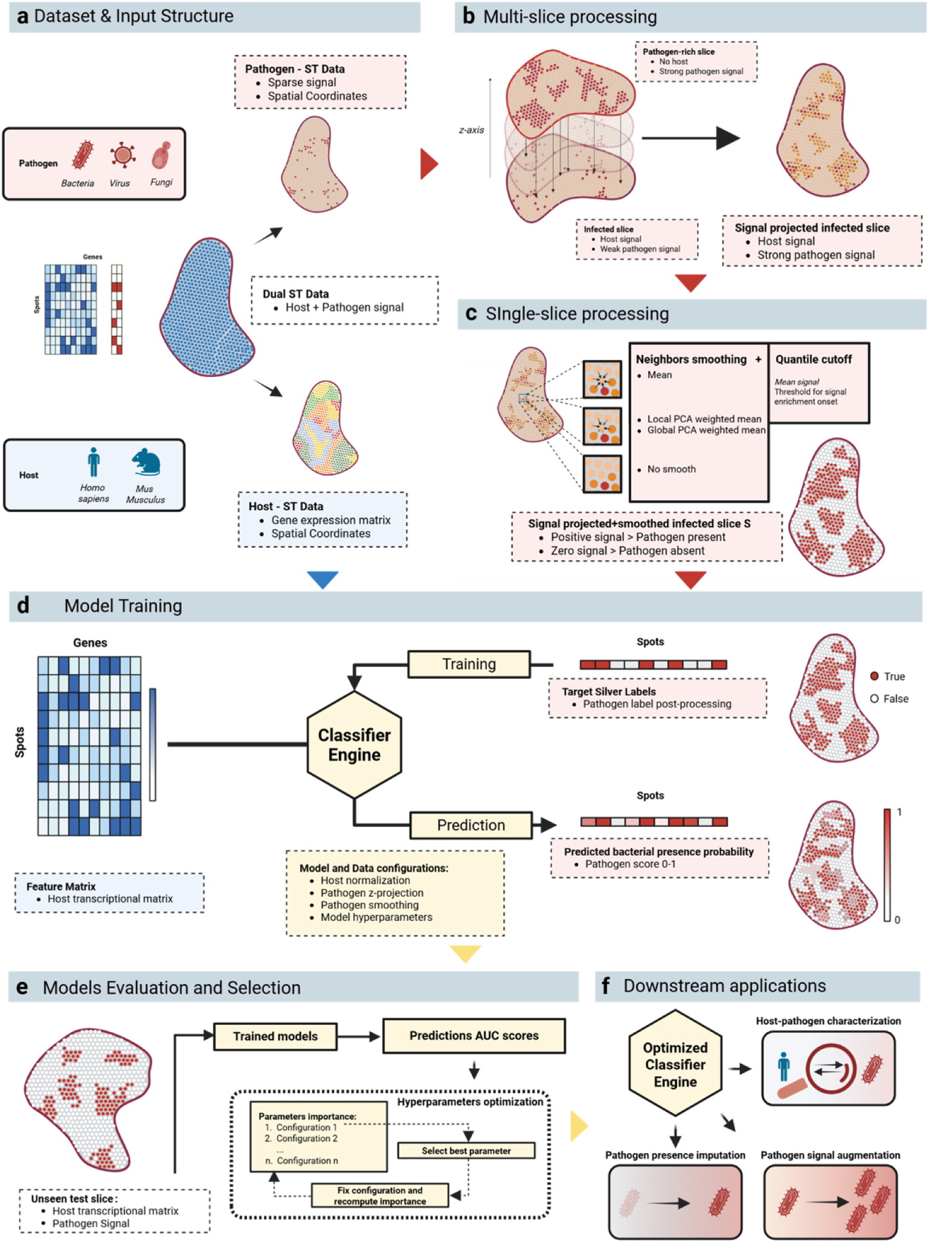
Overview of the PatHoST framework: (a) Input data structure: The framework accepts dual ST data containing both host and pathogen signals. The framework is pathogen-agnostic and cross-species adaptable. Host ST data is used as the feature matrix for model training, while sparse pathogen signals are directed to the silver label construction modules; (b) Multi-slice processing step: to enhance pathogen signal in infected slices, a Z-axis projection strategy is applied by combining data from adjacent pathogen slices. This step increases the detectable pathogen signal while maintaining spatial alignment with the host signal; (c) Single-slice processing step: Pathogen matrices are spatially smoothed to reduce noise and strengthen local signal continuity across neighboring spots. A binary label is derived by thresholding the smoothed pathogen signal, defining the silver labels for the subsequent model training step; (d) Training Step: Multiple model types and configurations are trained using host transcriptomic data as input and the silver binary label as target. The outputs of the models are continuous values between 0 and 1. The different trained models are passed to the evaluation step for final selection; (e) Model evaluation and Selection step: An iterative data-driven optimization strategy evaluates model configurations based on AUC scores from an unseen external test slice; (f) Downstream applications: The final optimized classifier enables downstream applications, including host-pathogen characterization, pathogen presence imputation, and signal augmentation in datasets where microbial capture is suboptimal or not performed.

Once trained, the optimized model can be applied to multiple downstream analyses, including the investigation of host–pathogen interactions, augmentation of sparse pathogen signal, and prediction of pathogen spatial distributions in tissues where pathogen transcripts are only partially detected or entirely unavailable (Figure 1f).

To evaluate the robustness and generalizability of the framework, PatHoST was assessed across two complementary experimental benchmarks. Case Study 1, based on the murine *Mycobacterium abscessus* dataset, exploits a uniquely designed experimental system incorporating pathogen-only donor sections to optimize the complete workflow, from pathogen signal enhancement to classifier selection, and to investigate the biological interpretability of the resulting predictions. Case Study 2, based on an independent human SARS-CoV-2 dataset, evaluates the transferability of the optimized framework to a distinct host–pathogen system characterized by substantially higher pathogen signal sparsity.

A summary of the experimental objectives, validation strategy, and evaluation metrics for each analysis is provided in Table 1.

**Table 1:** Roadmap of the experimental validation and benchmarking conducted. Column 1 reports the Case study index, Column 2 the Analysis step performed, Column 3 the Objective and Rationale of the experiment and Column 4 the major key metrics adopted for evaluation.

| # | Analysis Step | Objective & Rationale | Key Metrics |
| --- | --- | --- | --- |
| 1 | *Probe comparison* | Validate pathogen-only slices signal | Neg. Binomial Reg.,<br>Probe counts |
| 1 | *Signal Enhancement* | Evaluate Z-projection and smoothing impact for silver label generation. | Signal-to-noise ratio,<br>Quantile distribution |
| 1 | *Model Selection* | Identify robust classifier architecture and hyper-parameters/configuration selection. | AUC (External Test Set) |
| 1 | *Discretization* | Assess the optimal discretization algorithm for the prediction scores. | ARI, Silhouette, F1-score |
| 1 | *Biological Discovery* | Investigate PatHoST biological discoveries and comparison with external data | Spearman corr.,<br>GSEA, PBMC overlap |
| 2 | *Model Selection* | Identify robust classifier architecture and hyper-parameters/configuration selection. | AUC (External Test Set) |
| 2 | *Biological Discovery* | Investigate PatHoST biological discoveries | Spearman corr.,<br>GSEA |

### Experimental Design and Validation of Pathogen-only Slices

To investigate the technical determinants of incomplete pathogen detection in dualST, we evaluated two complementary experimental strategies aimed at improving bacterial transcript recovery: (i) enzymatic permeabilization of the bacterial cell wall using lysozyme to enhance probe accessibility and ligation efficiency, and (ii) the generation of pathogen-only tissue sections by removing host probes to reduce competition from highly abundant host transcripts. Pathogen transcript recovery was compared between conventional dualST sections (*S*_1_*_−_*_4_) and adjacent pathogen-only donor sections (*S_GT_* _1,2_), with and without lysozyme treatment (Figure 2a,b). Conventional dualST libraries detected pathogen transcripts in only a small fraction of the tissue, with pathogen-positive capturing spots ranging from 2.1% (*S*_1_) to 3.5% (*S*_4_) (Figure 2c,d). In contrast, pathogen-only donor sections substantially increased transcript recovery, reaching a maximum of 16.5% pathogen-positive capturing spots in *S_GT_* _1_. This increase was accompanied by a marked enrichment in pathogen-derived unique molecular identifiers (UMIs), supporting the presence of technical competition between host and pathogen probe capture. Beyond increasing transcript abundance, pathogen-only sections markedly extended the spatial organization of the detected pathogen signal. Whereas pathogen-positive spots in conventional dualST libraries appeared sparse and spatially fragmented, pathogen-only sections revealed contiguous high-density pathogen regions that more closely reflected the tissue bacterial loads and underlying tissue architecture (Figure 2a,b). Consistently, the Spatial Coherence Index, representing the percentage of positive spots with positive neighbors spots) increased from 40–51% in dualST sections to 67.4% in pathogen-only sections (Figure 2d). Similar improvements were observed for both targeted bacterial genes (*rpoB* and *lsr2*) (Figure 2e). To quantify the contribution of each experimental factor, we fitted a Negative Binomial regression model including sequencing batch, lysozyme treatment, and the presence or absence of host probes as explanatory variables. Removal of host probes was the only factor significantly associated with increased pathogen transcript recovery (Figure 2f), whereas neither lysozyme treatment nor sequencing batch significantly affected the number of recovered pathogen UMIs. Finally, we evaluated whether pathogen-positive spots detected in conventional dualST libraries represented genuine biological signal despite their sparse distribution. Following spatial registration with the corresponding pathogen-only donor sections (Supplementary Figure S1), approximately 50% of pathogen-positive spots directly overlapped high-density pathogen regions, while 74.6–87.3% were located within a single-neighborhood distance, increasing to 94.8–99.2% within an extended neighborhood (Figure 2g). These results demonstrate that pathogen-only sections preserve the underlying spatial organization of the infection while substantially increasing pathogen transcript recovery, thereby providing reliable high-density donor maps for multi-slice spatial projection and the generation of robust silver-standard labels within the PatHoST framework.

**Figure 2:**
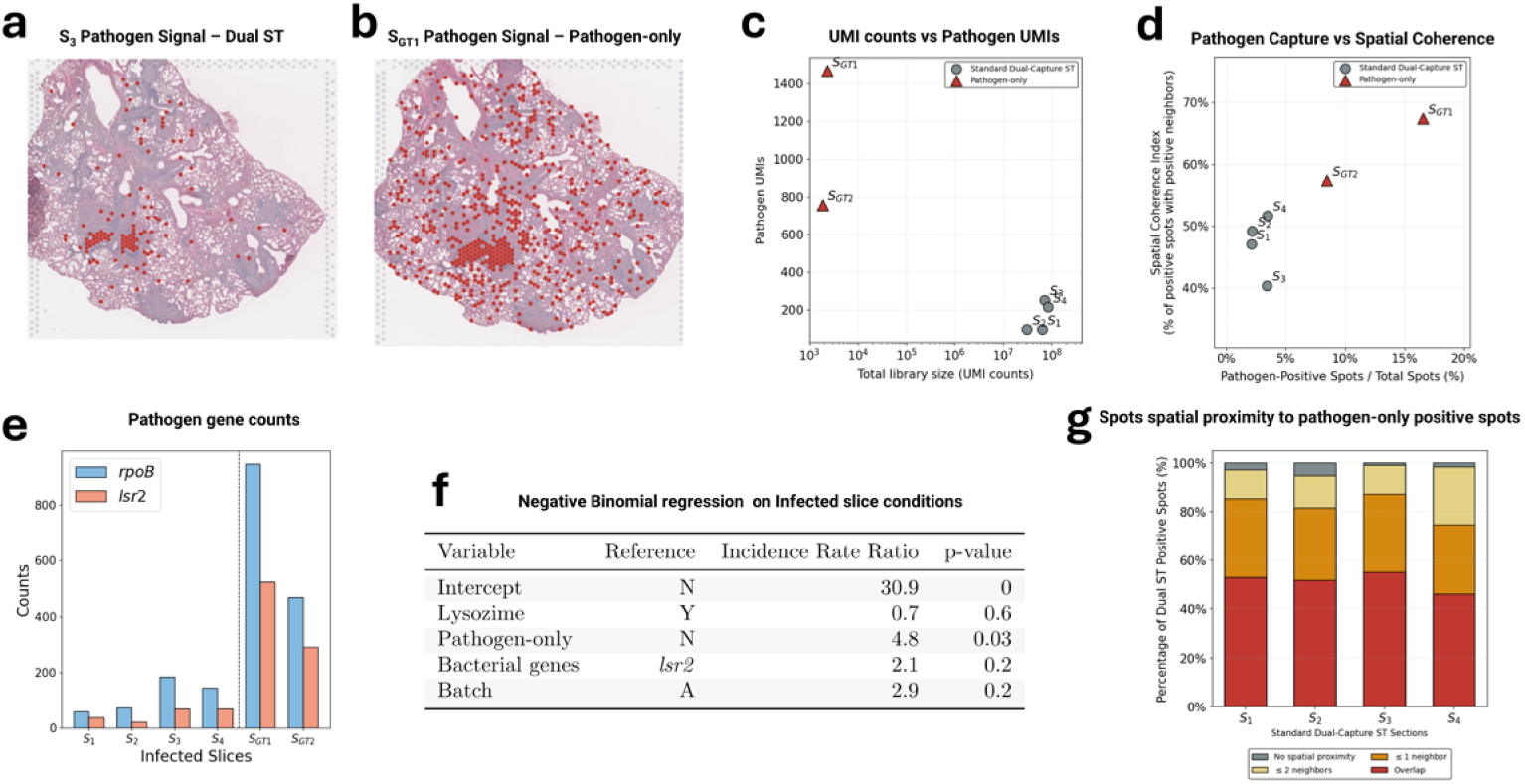
Host-depleted probe design rescues transcript density and spatial topography of sparse pathogen signals on Dataset 1 (murine lung chronically infected with *M. abscessus*. (a, b) Representative spatial distribution of pathogen-positive spots (≥ 1 UMI count, indicated by red dots) overlaid on H&E histological sections of infected murine lungs for standard Dual ST (*S*_3_) (a) and host-depleted pathogen-only (*S_GT_* _1_) (b) configurations. (c) Comparison of total library size and Pathogen UMIs across sections. (d) Spatial Coherence Analysis across all experimental sections. The scatter plot illustrates the relationship between the absolute percentage of pathogen-positive spots over total spots (x-axis) and the Spatial Coherence Index (y-axis, defined as the percentage of positive spots sharing a physical boundary with at least one adjacent positive spot). Standard dual-capture sections in grey circles and pathogen-only sections in red triangles. (e) Absolute UMI counts for target pathogen mRNAs (*rpoB* and *lsr2*) across standard dual-capture libraries (*S*_1_*_−_*_4_) and host-depleted sections (*S_GT_* _1_*_−_*_2_). (f) Negative Binomial regression modeling incidence rate ratios on infected slice conditions. (g) Spatial proximity analysis evaluating the distance of sparse Dual ST positive spots to the nearest pathogen-only niches. The cumulative bar plot quantifies the percentage of isolated Dual ST positive spots overlapping (red bars) or residing within a strict single-step neighborhood radius (orange bars) or a two-step extended local radius (yellow bars) of the aligned pathogen-only landmarks.

### Case Study 1: *Mycobacterium abscessus* chronic infection in murine lung

To comprehensively evaluate PatHoST, we assembled all spatial transcriptomic datasets generated from the *M. abscessus* chronic lung infection model, including both conventional dualST sections and the dedicated pathogen-only donor sections described above (Supplementary Table S1). We characterized the host trascriptional profiles, by performing integration for slices *S*_1_*_−_*_4_ and performing cluster analysis on the integrated dataset and on slice *S_EXT_* _1_ (Supplementary Figure S2). The availability of consecutive pathogen-only and dualST tissue sections enabled a systematic evaluation of every major component of the framework, including multi-slice Z-axis projection, signal enhancement, classifier optimization, and validation on an independent external dataset.

We systematically tested the implemented preprocessing techniques, data projection approaches, smoothing strategies, and model architectures. The tested configurations are summarized in Supplementary Table S2.

#### Signal projection and smoothing

To evaluate the contribution of spatial signal enhancement to latent pathogen signal recovery, we first assessed the projection and smoothing modules of PatHoST (Figure 3a-c). By spatially aligning consecutive infected tissue sections, including both conventional dualST and pathogen-only donor sections, the Z-axis projection substantially increased both the spatial coverage and the intensity of the detected pathogen signal. Subsequent spatial smoothing further reinforced local signal continuity while reducing stochastic sparsity across neighboring spots. The enhanced pathogen signal was subsequently binarized to generate the silver-standard labels used for classifier training (Figure 3a). To determine an appropriate threshold, we evaluated a range of quantile cutoffs across all smoothing strategies. Quantile thresholds ≥ 0.6 produced signal distributions that differed significantly from the unfiltered data (Supplementary Figure S3a), indicating that lower-intensity spots were predominantly associated with background noise. A cutoff of 0.7 was therefore selected as the optimal compromise between noise removal and retention of high-confidence pathogen-positive spots. Overall, each stage of the spatial enhancement workflow progressively increased the recoverable pathogen signal (Figure 3c). The proportion of pathogen-positive spots increased from approximately 3% in the raw data to 20% following Z-axis projection and reached nearly 80% after spatial smoothing. Application of the selected quantile threshold subsequently reduced the signal to approximately 30% of tissue spots, yielding the final silver-standard labels that combined expanded pathogen coverage with high spatial confidence for downstream model training.

**Figure 3:**
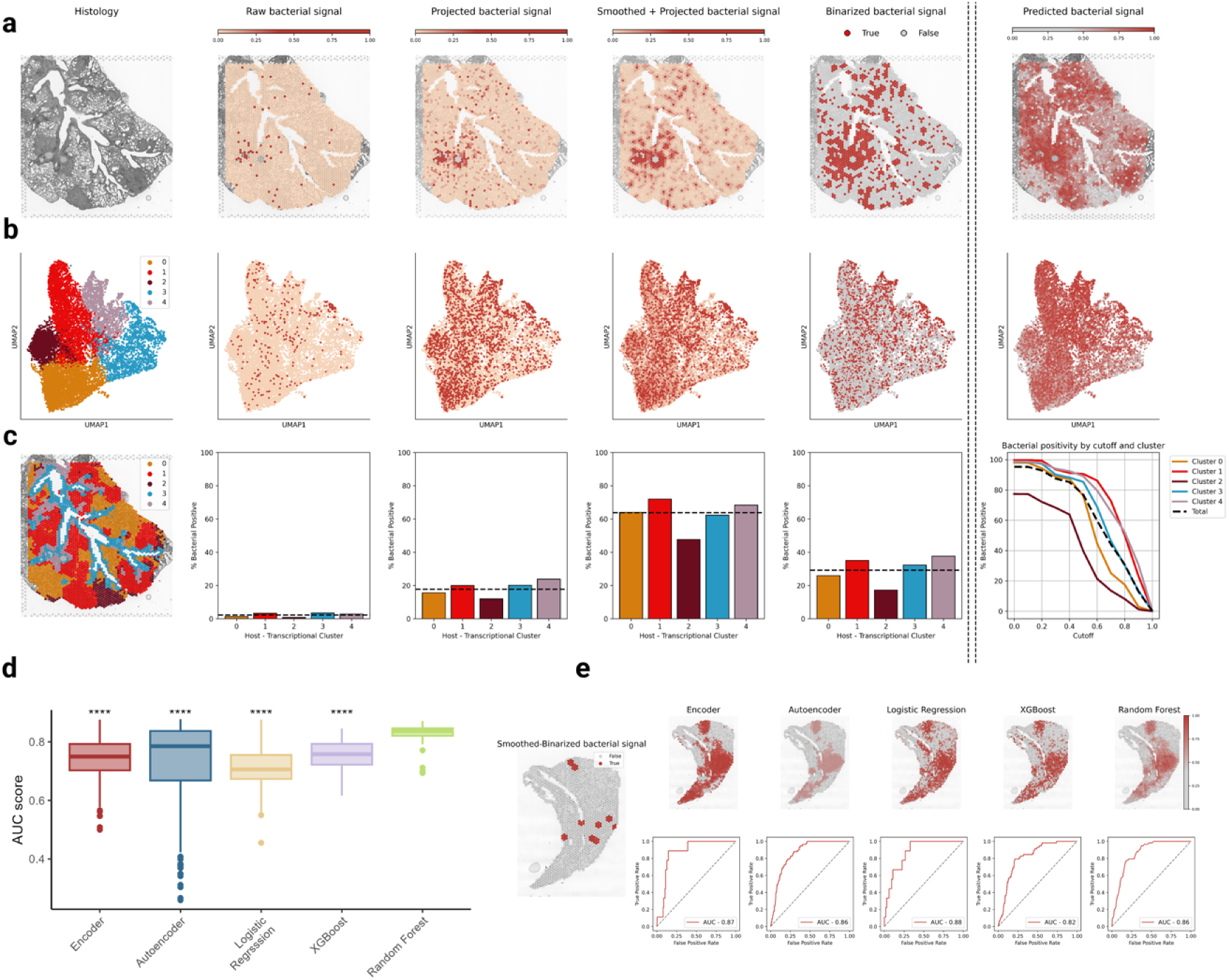
Performance evaluation and signal recovery on Dataset #1 (murine lung chronically infected with *M. abscessus*). (a) Overview of the spatial transformation pipeline applied to the example histological slice *S_1_*. The figure shows the histological section with the spatial transcriptomics spot grid; (b) UMAP low-dimensional representation of host transcriptomic clusters of slices *S*_1_*_−_*_4_ ; (c) summary statistics and cluster-level distributions. (a-c) From left to right, the panels illustrate the host transcriptomic clusters as reference, the raw bacterial signal, the bacterial signal projected from the pathogen-only slice *S_GT1_*, the projected signal after spatial smoothing using the Global PCA-Weighted Neighborhood Mean, the binarized signal used as labels for model training, and the prediction of a representative model expressed as continuous probabilities between 0 and 1. The bottom-right panel shows the variation in the percentage of bacteria-positive spots within each cluster across different binarization cutoffs, with colored lines representing individual clusters; (d) Distribution of AUC scores for the five model types across the explored hyperparameter space. Each model’s performance distribution was tested against the Random Forest distribution; (e) From left to right: bacterial signal of the external slice *S_EXT1_* after smoothing and binarization; prediction of bacterial signal by the five selected models expressed as continuous probabilities; and their respective ROC curves with associated AUC scores.

#### Performance assessment

To comprehensively evaluate PatHoST, we systematically explored the complete hyperparameter space across all implemented classifier architectures (Supplementary Table S2). Model performance was assessed on the independent external dataset *S_EXT_* _1_, an orthogonal dualST section generated from the same chronic *M. abscessus* infection model in a different animal, thereby providing a stringent evaluation of model robustness and generalizability. Before comparing classifier architectures, we quantified the specific contribution of the donor-derived Z-axis projection module. Models trained using only host gene expression achieved Area Under the Receiver Operating Characteristic Curve (AUC) values exceeding 0.80 (Supplementary Figure S3c), confirming that host transcriptional programs inherently reflect the spatial footprint of infection. However, incorporating pathogen information through multi-slice Z-axis projection consistently improved predictive performance, particularly on the external test dataset. These results demonstrate that donor-derived spatial projection expands the density and quality of the silver-standard labels, resulting in more robust decision boundaries and improved model generalization. Across the complete hyperparameter search, all classifier families achieved median AUC values above 0.70 on *S_EXT_* _1_ (Figure 3d). Random Forest models exhibited the highest median performance (*>* 0.80) together with the narrowest performance distribution, indicating superior robustness across different preprocessing and hyperparameter configurations. In contrast, the remaining classifier architectures showed significantly broader performance distributions and lower overall predictive accuracy. Stratification of model performance across individual hyperparameters further demonstrated that specific configurations consistently improved prediction accuracy, including dropout rates between 0.3 and 0.4 for neural-network architectures and the omission of pathogen count normalization (Supplementary Figure S3b).

The distributions obtained from the exhaustive hyperparameter search were subsequently used to identify the optimal configuration for each classifier through the iterative model selection procedure described in the Online Methods section ”Model selection” (Figure 1e). The resulting optimal models are summarized in Supplementary Table S3 and evaluated on the independent *S_EXT_* _1_ dataset in Figure 3e. Under their respective optimal configurations, Logistic Regression (*AUC* = 0.88) achieved the highest predictive performance, followed by the Encoder (*AUC* = 0.87), Autoencoder (*AUC* = 0.86), Random Forest (*AUC* = 0.86), and XGBoost (*AUC* = 0.82).

Finally, analysis of the iterative optimization procedure revealed that the relative importance of preprocessing choices and hyperparameters differed substantially across classifier architectures, indicating that no universally optimal data-processing strategy exists. Tree-based methods were primarily influenced by spatial preprocessing, particularly the choice of smoothing strategy and pathogen-only donor section used for Z-axis projection, whereas neural-network architectures were more strongly driven by internal hyperparameters such as dropout rate. Logistic Regression and XGBoost generally favored simpler host gene expression representations, while Random Forest derived greater benefit from spatial signal enhancement. Collectively, these findings demonstrate that although optimal configurations are model-specific, donor-derived spatial augmentation represents a consistent determinant of predictive performance across the PatHoST framework.

Notably, while the AUC shows robustly high values across silver labels and raw signal, the F1-score remains systematically low (0.04 on raw and 0.21 on silver labels; Supplementary Figure S3d) due to technical dropouts. This widespread suppression highlights how overlap-based metrics are severely bottlenecked by spatial sparsity, validating our choice to evaluate performances through AUC.

#### Defining Confidence Zones via Thresholding

While the predicted likelihood of pathogen presence provides the most informative representation for quantitative downstream analyses, practical applications often benefit from the identification of discrete confidence regions. We therefore evaluated multiple data-driven strategies to discretize the continuous prediction scores, including median-based thresholding, hierarchical clustering (k = 2 and k = 3), fuzzy clustering (k = 3), and k-means clustering (k = 2 and k = 3). Among the evaluated approaches, both k-means solutions performed favorably across multiple evaluation metrics (Supplementary Figure S4a). However, the three-cluster (k = 3) solution provided the best compromise between inter-cluster separation, measured by the Calinski–Harabasz index, and agreement with the binarized silver-standard labels, assessed by the F1 score. Moreover, the additional intermediate class was particularly suited to the objective of PatHoST, namely the recovery of latent pathogen signal beyond the experimentally detected pathogen-positive spots. We therefore selected the three-cluster k-means solution for all downstream analyses. The selected model partitions the continuous prediction scores into three spatially coherent confidence regions—low, medium, and high (Supplementary Figure S4b-d). The high-confidence region closely overlapped the experimentally observed pathogen signal, whereas the medium-confidence region predominantly represented pathogen-positive areas inferred by PatHoST, thereby extending the spatial landscape of infection while preserving biological plausibility. The resulting confidence regions also showed clear correspondence with host tissue organization. High-confidence regions were predominantly associated with inflammatory (Cluster 1) and submucosal (Cluster 4) transcriptional domains, whereas parenchymal and airway regions (Clusters 0 and 3) were preferentially enriched for medium-confidence predictions (Supplementary Figure S2a-c and Supplementary Figure S4e-g). Together, these results demonstrate that discretization preserves the biological structure of the continuous prediction landscape while providing an intuitive framework for downstream spatial interpretation.

#### Biological interpretability of PatHoST predictions

To investigate the biological relevance of PatHoST predictions, we correlated the predicted likelihood of pathogen presence in the independent dataset *S_EXT_* _1_ with host gene expression. The top-ranked genes identified across the four classifier architectures showed strong concordance and were consistently associated with immune and inflammatory pathways (Figure 4a). Moreover, these four groups of genes could be associated to different areas of the tissue slice, providing a complementary view to host-based transcriptional clustering (Figure 4b-c and Supplementary Figure S2d-e). For example, genes involved in MHC-II signaling and T-cell recruitment were preferentially enriched within localized regions of transcriptional Cluster 1, corresponding to focal inflammatory niches (Supplementary Figure S2d-e). To determine whether the inferred pathogen signal reflected genuine host biology rather than computational artifacts, we evaluated an independent host transcriptional signature previously derived exclusively from raw bacterial probe counts, without signal enhancement, in two of the training sections (*S*_1,2_) [2]. Projection of this signature onto the independent *S_EXT_* _1_ dataset closely matched the spatial distribution predicted by PatHoST (Figure 4d-f), indicating that latent pathogen signal recovery preserves bona fide infection-associated host transcriptional programs. Consistent with this observation, the medium-confidence region identified by PatHoST displayed a significant enrichment of the infection signature compared with the low-confidence background (*P_value_ <* 0.01; Figure 4g), further supporting the biological validity of the discretization strategy. We next investigated whether PatHoST could resolve spatially distinct regulatory programs associated with different infection niches. Using the Random Forest model as a representative example, we correlated transcription factor (TF) expression (AnimalTFDB [10]) with the predicted likelihood of pathogen presence separately within Cluster 0 (parenchyma) and Cluster 1 (inflamed tissue). TFs exhibiting a correlation coefficient *>* 0.1 and an inter-cluster difference *>* 0.2 were selected for downstream analysis (Figure 4h). This approach identified two distinct regulatory landscapes whose TF expression positively tracked the local pathogen prediction score (Figure 4i). Cluster 0 was characterized by TFs associated with adaptive immunity and host defense against mycobacterial infection, including Nfkb2, Irf8, and Trp53 [11, 12]. In contrast, Cluster 1 was enriched for TFs linked to granuloma-associated responses, cellular stress adaptation, nutritional immunity, and iron sequestration, including Atf5, Snai2, and Ltf [13–15]. Notably, many of these TFs were not identified by conventional clustermarker analysis and were, in several cases, globally downregulated relative to the remainder of the tissue. Their positive spatial correlation with PatHoST predictions therefore indicates that they represent localized pathogen-associated regulatory activity rather than general markers of cluster identity. Collectively, these analyses demonstrate that PatHoST captures the spatial heterogeneity of host–pathogen interactions beyond conventional transcriptomic analyses. To further assess the clinical relevance of these findings, we examined the expression of cluster-specific and shared TFs in an independent bulk RNA-seq dataset of peripheral blood mononuclear cells from patients with *M. abscessus* pulmonary disease [16]. All three TF sets were significantly upregulated in infected patients relative to non-infected controls (Figure 4j), supporting the biological and translational relevance of the spatial regulatory programs identified by PatHoST.

**Figure 4:**
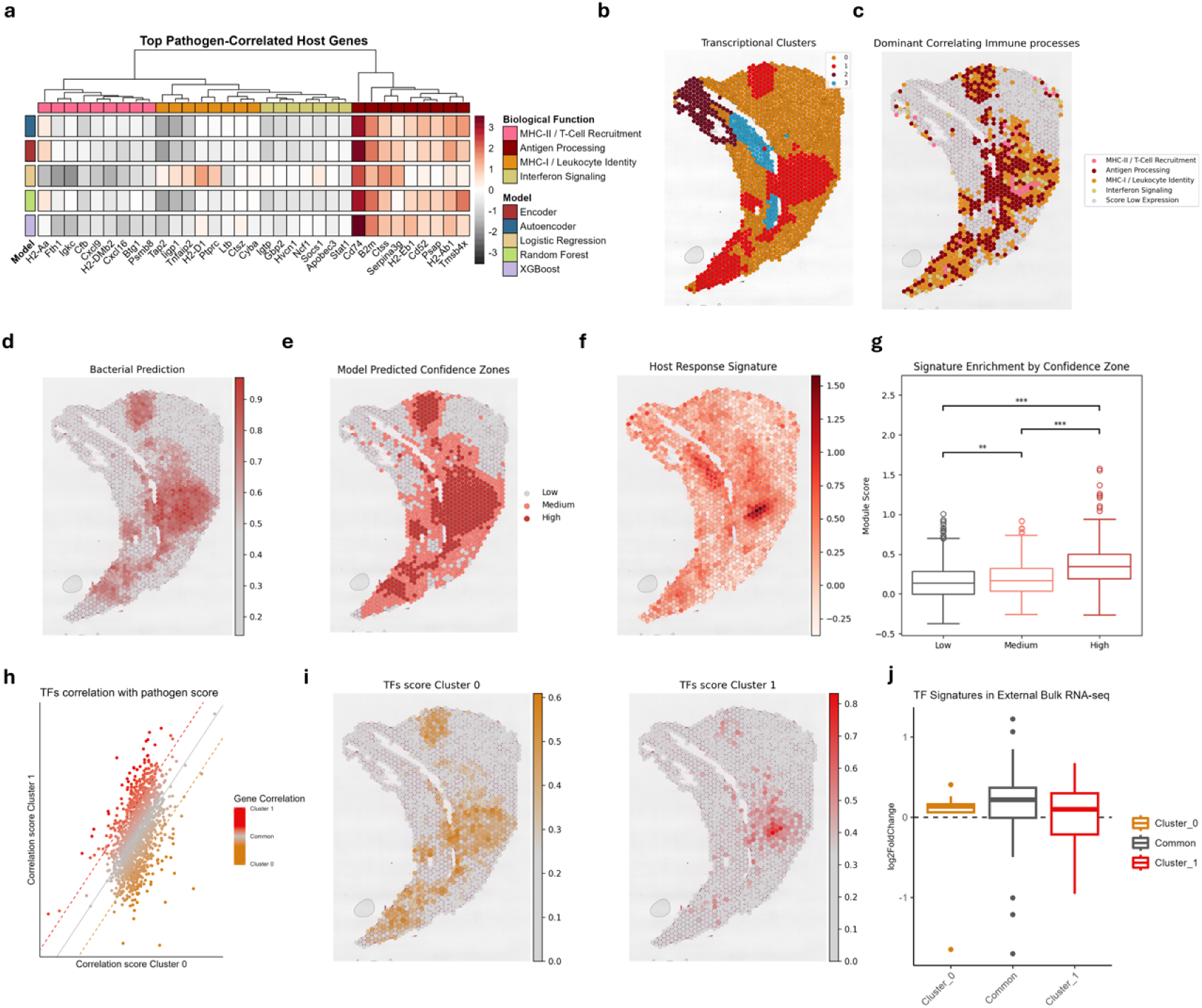
Biological evaluation for PatHoST predictions on Dataset #1 (murine lung chronically infected with *M. abscessus*). (a) Heatmap illustrating the top 20 host genes most strongly correlated with the prediction score across the Framework models, with hierarchical clustering to identify gene programs conserved among the models; (b) Spatial map of host transcriptional clusters identified in *S_EXT1_*; (c) Spatial map of *S_EXT1_*, indicating the highest expressed gene module score (derived from correlating genes) for each spot, illustrating the tissue-level distribution of the predicted biological response; (d) Spatial map of *S_EXT1_* showing the continuous prediction score (0-1) generated by PatHoST; (e) Spatial map of *S_EXT1_* showing the predicted confidence zones (Low, Medium, High) using a k-means approach for discretization with *k* = 3; (f) Spatial map of *S_EXT1_*, for the module score of Host response signature to *Mabs* derived from *Di Marco & Nicola et al.* [2]; (g) Boxplot of the host signature module score across the three model-predicted confidence zones; (h) Scatterplot comparing the Spearman correlation of host Transcription Factors (TFs) with the pathogen prediction score in Cluster 0 (Parenchyma) versus Cluster 1 (Focal Inflammation), highlighting TFs driving cluster-specific responses; (i) Spatial maps of *S_EXT1_* showing the gene module scores of the Transcriptional Factors associated with Cluster 0 by correlation with the pathogen prediction score (left) and Transcriptional Factors associated with Cluster 1 (right); (j) Boxplots showing the log_2_ Fold Change (log_2_ FC) distribution in an external bulk RNA-seq cohort (PBMCs from *M. abscessus* patients vs. healthy controls) for the sets of TFs specific to Cluster 0 (orange), Cluster 1 (red), and shared (grey), supporting the clinical relevance of the spatially-derived regulatory signatures.

### Case Study 2: Generalization to SARS-CoV-2 infection in human lung

To evaluate the transferability of PatHoST beyond the experimental setting used for model development, we applied the framework to an independent ST dataset from human lungs infected with SARS-CoV-2 [3]. This dataset represents a substantially different biological context, involving both a different host species (human) and a distinct pathogen class (virus), thereby providing a stringent assessment of the framework’s generalizability. Healthy human lung sections from Mayr et al. [17] were included as controls (Supplementary Table S4). Because no adjacent donor sections were available, the Z-axis projection module was omitted while all remaining preprocessing and model-selection strategies described in Supplementary Table S2 were retained. Predictive models were trained using host transcriptomes from both infected and healthy donor sections and evaluated on an independent infected tissue section. The optimal models (Supplementary Table S5) achieved AUC values ranging from 0.61 (Autoencoder) to 0.78 (Random Forest) on the external test dataset (Figure 5a,b). Although predictive performance was lower than that observed in the *M. abscessus* case study, this decrease closely mirrored the reduction previously observed when Z-axis projection was removed from the bacterial analysis, supporting the conclusion that donor-derived spatial augmentation substantially enhances performance while the core PatHoST framework remains transferable across different host–pathogen systems.

To investigate the biological relevance of these predictions, we performed host transcriptional clustering of the human lung tissue, identifying four major tissue domains: Cluster 0 (Parenchyma), Cluster 1 (Reactive Adventitial Niche), Cluster 2 (Quiescent Alveolar Stroma), and Cluster 3 (Inflammatory Myeloid Core) (Figure 5c). Correlation analysis between TF expression and the predicted likelihood of pathogen presence revealed distinct regulatory programs associated with viral localization in each tissue niche. Cluster 1 was characterized by TFs involved in acute inflammation (JUNB), hypoxia (HIF1A), and immune exhaustion (FOXO1) [18–20]. Cluster 0 was enriched for regulators associated with tissue damage and vascular remodeling (TP53, NR2F2) [21, 22], whereas Cluster 3 showed enrichment for transcriptional regulators linked to pulmonary fibrosis (SMAD3, SMAD4, ZEB2) [7, 23] (Figure 5d). Even after spatial smoothing, the experimentally detected viral signal remained sparse within the external test section (Figure 5e). In contrast, PatHoST reconstructed a continuous spatial landscape of predicted viral localization (Figure 5f), assigning the highest prediction confidence to the Reactive Adventitial Niche (Cluster 1) while simultaneously identifying additional lower-confidence viral-associated regions distributed throughout the tissue (Figure 5g). These findings demonstrate that PatHoST generalizes across distinct host species, pathogen classes, and experimental datasets while preserving biologically meaningful spatial regulatory programs.

**Figure 5:**
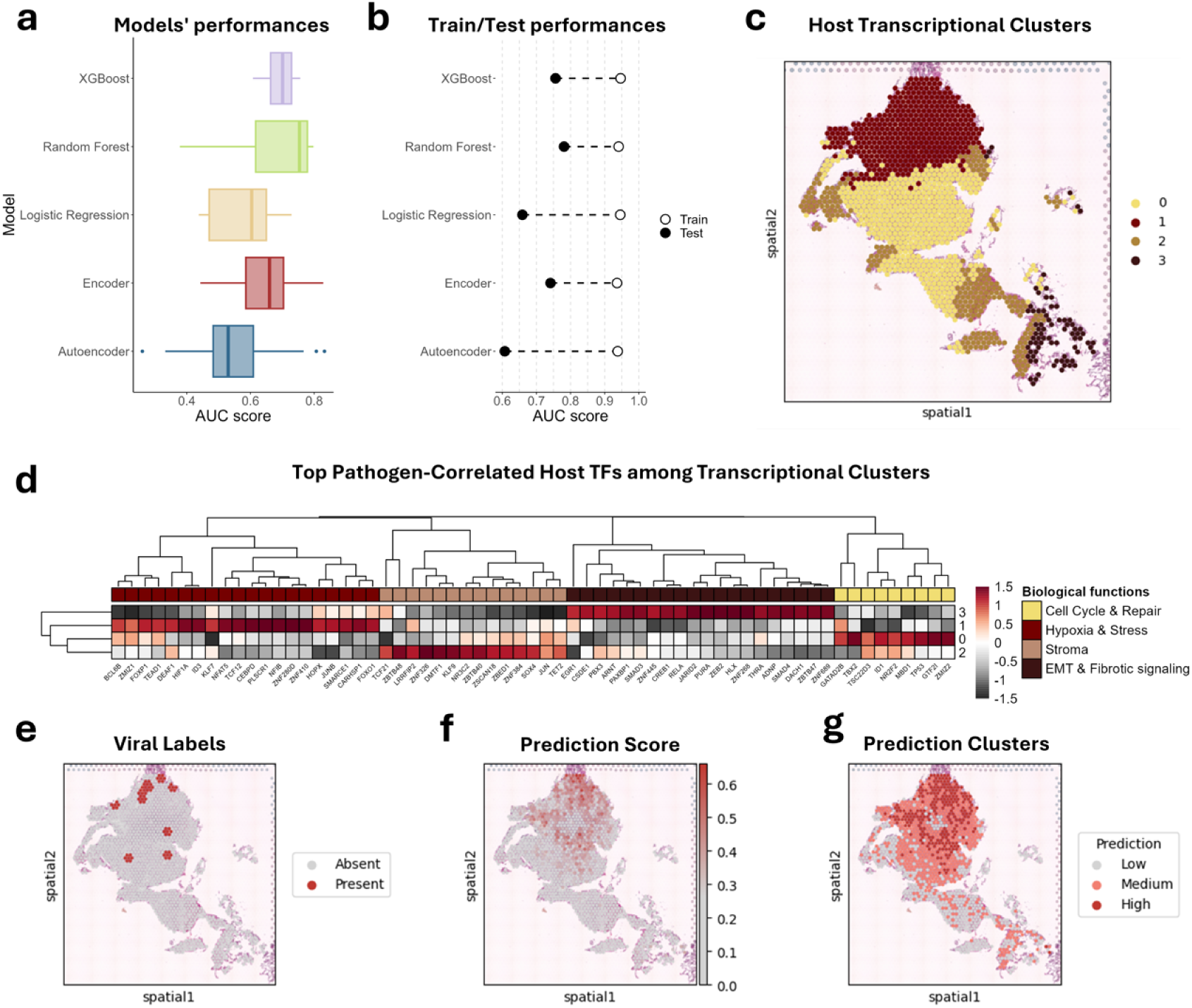
Performance assessment and biological evaluation for PatHoST predictions on test slice *C*_4_ from Dataset #2 (human lung with SARS-CoV-2). (a) Distribution of AUC scores for the five model types across the explored hyperparameter space; (b) Dot plot reporting the drop in accuracy performance for each model type between train and test datasets; (c) Spatial map of host transcriptional clusters identified in the test slice; (d) Heatmap illustrating the top 20 host genes most strongly correlated with the Random Forest prediction scores across the Host transcriptional clusters; (e) Spatial map of the viral labels after the Global PCA-Weighted Neighborhood Mean smoothing process; (f) Spatial map of the viral prediction score from the Random Forest model; (g) Spatial map of the Clustering of viral prediction scores from the Random Forest model into confidence categories (Low, Medium, High).

## Discussion

In this study, we presented PatHoST, a computational framework designed to overcome the limited pathogen signals quantification in ST. To date, efforts have largely focused on enhancing direct capture technologies or alignment algorithms; while current ST technologies offer high resolution for host tissues, microbial transcripts often fall below detection limits or appear stochastically distributed, hindering the reconstruction of infection maps. Our results demonstrate that the host transcriptional landscape retains a robust ”echo” of pathogen presence, which can be decoded using machine learning even when the direct pathogen signal is elusive.

A central contribution of this work is the conceptualization of pathogen detection in ST as a weakly supervised learning task. Traditional supervised learning relies on ”gold standard” labels perfectly accurate annotations of the target variable. In the context of infected tissue sections, obtaining such ground truth is challenging: direct pathogen sequencing is limited, and aligning adjacent slides introduces spatial noise (label uncertainty). PatHoST addresses this by generating ”silver labels” via our Z-projection and spatial smoothing pipeline. We define this approach as weakly supervised because the training targets are inferred and enhanced rather than directly observed with absolute precision. By training models on these noisy but signal-rich labels, we force the algorithms to learn the robust, underlying correlations between host gene expression and pathogen presence, effectively filtering out the stochastic noise inherent in the labeling process. The capability of our models to generalize to external datasets (*S_EXT_* _1_) confirms that the models successfully learned the biological signature of infection rather than overfitting the artifacts of the projection technique.

Our comparative analysis highlighted the improvement added by the Z-projection strategy, especially in the presence of pathogen-rich slides, suggesting that strategies to improve bacterial detection are required. Models trained solely on the sparse signal of the infected slice are less capable of generalizing, achieving lower performance on external validation sets. By projecting the signal using Z-projection, we boosted the signal-to-noise ratio, providing the models with a sufficient volume of positive training examples. While this might introduce a degree of spatial mismatch, our results indicate that the benefit of increased label density outweighs the cost of spatial imprecision, especially when robust smoothing strategies (like Global PCA-Weighted Mean) are applied to enforce local coherence.

A notable observation is the divergence between F1 scores and AUC metrics when evaluated against Raw Counts. While PatHoST consistently achieved high discriminative performance (AUC *>* 0.80), the F1 scores were significantly lower across all tested methods. This discrepancy reflects the incompleteness of the observed pathogen signal rather than reduced predictive performance due to the mismatch between latent signal recovery and technical sparsity inherent to this technology [24]. Because raw counts contain only a limited number of detected pathogen-positive spots, predictions extending beyond these observations are penalized as false positives, even when they likely correspond to true infection regions. In contrast, the consistently high AUC indicates that PatHoST correctly ranks pathogen-associated locations despite the incompleteness of the reference labels. These results suggest that AUC provides a more informative measure of performance than overlap-based metrics such as F1 when evaluating pathogen reconstruction from sparse observations. Beyond predictive performance, the validity of a weakly supervised approach ultimately lies in its biological interpretability. PatHoST identified distinct transcriptional programs associated with the predicted pathogen distribution, including adaptive immune markers (Nfkb2, Irf8) in the parenchyma and stress- and granuloma-associated markers (Atf5, Ltf) in inflamed regions. Importantly, several of these regulators were not detected by conventional cluster-level differential expression analysis but emerged when gene expression was correlated with the spatial predictions generated by the model. This suggests that PatHoST captures local transcriptional heterogeneity that may be overlooked by standard clustering approaches. Consistent with this observation, a host signature previously associated with *M. abscessus* infection [2] was enriched within our predicted ’Medium’ confidence zone, supporting the biological relevance of the reconstructed pathogen distribution.

Finally, the robustness of the framework was evidenced by its application to a completely independent dataset: human lung tissue infected with SARS-CoV-2. Despite the fundamental biological differences between a bacterial (extracellular/intracellular mycobacterium) and a viral infection, and the shift from a murine to a human host, PatHoST successfully reconstructed viral distributions and identified relevant hypoxia and inflammation signatures (HIF1A, JUNB). This suggests that the core principle of our weakly supervised framework, leveraging the host’s reaction as a proxy for pathogen presence, is broadly applicable across infectious disease biology.

Our approach is not without limitations. Z-projection requires the availability of adjacent tissue sections, which may not always be feasible in clinical retrospective studies. Furthermore, the resolution of our predictions is inherently bound by the diffusion of host signaling; while we can predict the niche of infection, pinpointing individual bacteria remains beyond the scope of host-inferred signals. Future iterations of this framework could incorporate multi-modal data or more advanced probabilistic graphical models to better quantify the uncertainty in the ”weak” labels, potentially refining the spatial resolution of the predictions.

Collectively, our findings demonstrate that host transcriptomes contain sufficient spatial information to improve the reconstruction of pathogen localization even when direct pathogen detection is incomplete. By framing pathogen reconstruction as a weakly supervised learning problem, PatHoST establishes a general computational strategy for integrating host and pathogen data in ST while enabling the discovery of localized host regulatory programs associated with infection. More broadly, our results suggest that the host transcriptome itself can serve as a spatial sensor of infection, providing a new framework for investigating host–pathogen interactions directly within intact tissues as dual spatial transcriptomics technologies continue to mature.

## Online Methods

### PatHoST Framework

PatHoST (Pathogen-Host Spatial Transcriptomics) is a modular computational framework for latent pathogen signal recovery from ST. By formulating pathogen detection as a weakly supervised learning problem, PatHoST exploits the host transcriptional response to infer pathogen spatial distributions when direct pathogen detection is incomplete. The framework is pathogen-agnostic readily applicable across different host species, pathogens and ST platform.

The workflow consists of four sequential modules (Figure 1) (A) multi-slice processing, in which consecutive tissue sections are aligned to enhance pathogen signal through spatial projection; (B) single-slice processing, where smoothing techniques reinforces signal coherence; (C) silver standard labels generation, which transforms the augmented pathogen signal into training labels; and (D) supervised learning, where machine learning models use host gene expression to predict the pathogen spatial distribution. The modular design of PatHoST enables flexible combinations of preprocessing, normalization and spatial signal enhancement strategies. Regardless of the selected configuration, the framework follows the same conceptual workflow: sparse pathogen observations are transformed into silver standard labels, which are subsequently used to train predictive models based exclusively on the host transcriptional response.

#### Input Data and Mathematical Representation

PatHoST operates on host-pathogen dualST datasets represented as transcriptional matrices. Given a tissue section, let *N* = {1, 2*, …, n*} denote the set of capturing spots and *G* = {1, 2*, …, g*} the set of detected genes. After preprocessing, gene expression is represented by the transcriptional matrix *X* ∈ ℝ*^n×g^*, where each row corresponds to a capturing spot, and each column to a gene.

The element *x_ij_* denotes the UMI count of gene *j* detected at capturing spot *i*. Each capturing spot is with two-dimensional spatial coordinates stored in the 2D capturing grid reported in matrix *C* ∈ ℝ*^n×^*^2^. The transcriptional matrix is subsequently partitioned into the host expression matrix *X_H_*, containing the set of host genes *G_H_*, and the pathogen expression matrix *X_P_*, containing the set of Pathogen genes *G_P_* (Figure 1a).

#### Algorithmic Steps and Modular Components

##### Step A: Multi-slice processing

When consecutive tissue sections are available, PatHoST exploits their spatial continuity to improve latent pathogen signal recovery by projecting pathogen observations across adjiacent slides (Figure 1b). This step increase the density of silver-standard labels while preserving the underlying tissue architecture. To align a donor section *S_j_* to a section *S_i_*, a set of *m* corresponding anatomical landmarks is manually identified on both sections. Landmark coordinates are centered by subtracting their respective centroids, after which the optimal rigid transformation is estimated using Singular Value Decomposition to minimize the distance between corresponding landmarks. The resulting rotation matrix is then applied to the spatial coordinates of *S_j_*, aligning the donor section to section *S_i_*.

Following alignment, pairwise Euclidean distances between capturing spots in the two sections are computed. For each spot in *S_i_*, the nearest neighbor in the transformed *S_j_* is identified, and only pairs separated by less than a user-defined distance threshold are retained as overlapping locations.

For each overlapping spot, pathogen expression is updated by combining the observed pathogen counts in the reference section with the corresponding signal projected from the donor section:

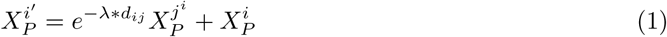

where *d_ij_* denotes the distance between sections in the z-stack (defined as the number of intervening section plus one), *λ* is the distance decaying coefficient and *X_P_* represents the pathogen expression matrix of the donor section. When multiple adjacent sections are available, the procedure is applied iteratively to integrate pathogen information across all donor sections.

##### Step B: Single-slice processing

To further improve latent pathogen signal recovery, PatHoST applies spatial signal enhancement to the pathogen expression matrix within individual tissue sections (Figure 1c). This module exploits spatial proximity and, optionally, host gene expression similarity to reinforce local pathogen signal coherence before silver-standard label generation.

PatHoST implements three naive spatial smoothing strategies: Neighborhood Mean, Local PCA-Weighted Neighborhood Mean, and Global PCA-Weighted Neighborhood Mean [25]. In addition, the framework is designed to integrate external imputation methods, allowing alternative signal enhancement strategies to be incorporated without modifying downstream analyses.

- Neighborhood Mean

The Neighborhood Mean provides a baseline spatial smoothing strategy based solely on physical proximity. For each capturing spot, pathogen expression is estimated as the average pathogen signal across its first-order spatial neighborhood.

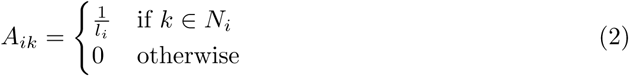

Where *N_i_* denotes the set of spatial neighbors of spot *i* (*i* included) and *l_i_* = |*N_i_*| represents the total number of spots in this set. The imputed pathogen transcriptional matrix is computed as *X^′^_P_* = *AX_P_*.

- Local PCA-Weighted Neighborhood Mean

The Local PCA-Weighted approach incorporates host gene expression to refine spatial smoothing. For each capturing spot, a principal component analysis (PCA) is computed using the spot and its immediate spatial neighbors. The resulting embedding (*L^i^* ∈ ℝ^(*l*^*^i^*^)^*^×^*^min(*l*^*^i,^*^50)^) is used to calculate the cosine similarities between neighboring spots, yielding a similarity matrix that favors transcriptionally similar neighborhoods:

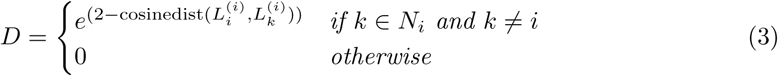

The similarity matrix is then normalized to produce the weighted adjacency matrix *A* ∈ ℝ*^n×n^*:

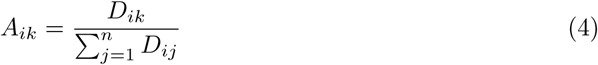

The imputed pathogen transcriptional matrix is calculated as:

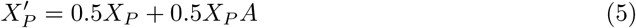

- Global PCA-Weighted Neighborhood Mean:

The Global PCA-Weighted strategy follows the same procedure but computes a single PCA is computed across the entire tissue section. Similarity weights are therefore derived from the global host transcriptional landscape rather than from local neighborhoods, enabling signal reinforcement over broader tissue domains.

Rather than representing alternative smoothing algorithms, these strategies capture different assumptions about tissue organization. Neighborhood Mean assumes that pathogen abundance varies primarily as a function of spatial proximity. In contrast, the PCA-weighted approaches additionally exploit host gene expression, under the assumption that transcriptionally similar spots are more likely to share similar pathogen states. The Local PCA emphasizes microenvironmental heterogeneity and sharp tissue transitions, whereas the Global PCA captures broader transcriptional gradients and provides greater robustness when local variation is dominated by technical noise

- Integration of external signal enhancement methods:

To demonstrate the modularity of the framework, we additionally implemented the MAGIC algorithm [6] as an alternative signal enhancement strategy. In this configuration, pathogen expression is reconstructed using the host gene expression manifold, illustrating how existing transcript imputation methods can be seamlessly incorporated into the PatHoST workflow.

#### Step C: Silver Labels Generation

Following signal enhancement (Steps A and B), pathogen expression is aggregated across all targeted pathogen genes for each capturing spot. Binary silver-standard labels are then generated using a data-driven quantile thresholding strategy to partition the tissue into pathogen-positive and pathogen-negative regions. Candidate quantile thresholds are evaluated using independent two-sample t-tests to identify informative silver-standard labels for downstream model training. The resulting binary labels are subsequently used as training targets for supervised learning.

These annotations are referred to as silver-standard labels because technical dropouts and incomplete pathogen detection preclude the definition of an error-free gold standard in dualST datasets. Rather than treating the observed pathogen counts as absolute ground truth, PatHoST formulates pathogen prediction as a weakly supervised learning problem, using the silver-standard labels as proxy annotations for latent pathogen signal recovery.

#### Step D: Classification

In the final stage, the silver-standard labels generated in Step C are used to train supervised models that predict the pathogen spatial distribution from host gene expression (Figure 1d). PatHoST supports multiple classification backends, including Logistic Regression and Random Forest implemented through scikit-learn [26], XGBoost [27], and encoder- and autoencoder-based classifiers implemented in pytorch [28]. The modular architecture also enables the integration of user-defined models.

The encoder and autoencoder classifiers share the same encoder and classification modules. The autoencoder architecture additionally includes a decoder trained to reconstruct the input host gene expression profile, thereby combining the classification objective with a reconstruction objective. Further architectural and training details are provided in the Supplementary Methods.

#### Probabilistic and Discrete Output

All classification models implemented in PatHoST produce continuous probability scores ranging from 0 to 1, representing the predicted likelihood of pathogen presence at each capturing spot. For downstream visualization and biological interpretation, these probabilistic predictions can be discretized using data-driven unsupervised clustering methods (e.g., k-means), enabling the identification of spatial regions with distinct levels of predicted pathogen burden.

#### Training

Supervised models are trained using the silver-standard labels generated in Step C as binary targets and the corresponding host gene expression profiles as input features. To mitigate the inherent class imbalance between pathogen-positive and pathogen-negative spots, the majority class was randomly under-sampled prior to model training. For conventional machine learning models (Logistic Regression, Random Forest, and XGBoost), the data were partitioned into training and independent test sets using an 80:20 split. For neural network-based classifiers (Encoder and Autoencoder), a three-way split comprising 64% training, 16% validation, and 20% independent test data was adopted to enable hyperparameter optimization and reduce overfitting. To evaluate model generalizability, trained classifiers can be applied to completely independent ST datasets that were not used during model training or hyperparameter optimization. This evaluation assesses the ability of PatHoST to estimate the predicted likelihood of pathogen presence across distinct biological samples and experimental conditions, providing a measure of robustness to biological variability and technical batch effects.

#### Model selection

To identify the optimal combination of preprocessing, signal enhancement, and classification parameters, PatHoST implements an iterative model selection strategy (Figure 1e). Model performance was primarily evaluated according to the ability to rank the predicted likelihood of pathogen presence, quantified using the AUC. Precision-based metrics, including the F1-score and Precision-Recall Area Under the Curve, were not used as primary optimization criteria because pathogen observations in dualST constitute a weakly supervised labeling scenario [9]. Due to incomplete pathogen detection, spots lacking observed pathogen signal cannot be assumed to represent true negatives, causing precision-based metrics to systematically underestimate predictive performance when biologically meaningful pathogen regions are recovered beyond the observed signal. Model selection was performed through an iterative optimization procedure. First, the AUC was computed for every evaluated pipeline configuration. A Random Forest regressor was then trained to model the relationship between pipeline components (e.g., normalization strategy, signal enhancement method, projection settings, and classifier hyperparameters) and the corresponding AUC scores. Feature importance analysis was subsequently used to identify the pipeline component with the greatest influence on predictive performance. This component was fixed to its best-performing value, defined by the highest median AUC across all evaluated configurations, and the remaining parameter space was re-evaluated. The procedure was repeated iteratively until a single optimized configuration was identified for each classifier architecture. This strategy enables PatHoST to adapt its configuration to the characteristics of individual datasets while reducing the risk of overfitting during model selection.

### Evaluation of discretizaton strategies

To convert the continuous predicted likelihood of pathogen presence into biologically interpretable spatial regions, PatHoST evaluates multiple discretization strategies applied to the predicted probability scores of each capturing spot. The evaluated approaches include median-based thresholding, Hierarchical Clustering, Fuzzy C-Means, and k-means clustering. The resulting partitions are assessed using complementary criteria that evaluate cluster quality, robustness, and biological consistency. Internal validity: cluster compactness and separation are quantified using the Silhouette Score and the Calinski–Harabasz Index, computed using their implementations in scikit-learn [26]. Algorithmic stability: the robustness of each discretization strategy is evaluated using bootstrap resampling (100 iterations, 80% random sampling). Partition stability is quantified by comparing the resulting cluster assignments with those obtained from the original data using the Adjusted Rand Index. External consistency: the biological relevance of the discretized regions is assessed by comparing the high-confidence region (defined as the cluster with the highest mean predicted probability) with the corresponding silver-standard labels using the F1-score.

### Biological interpretation of model predictions

To assess the biological relevance of PatHoST predictions, we investigated the relationship between the predicted likelihood of pathogen presence and host gene expression across spatial transcriptomic spots. Given a trained classifier, Spearman correlation coefficients were computed between the predicted probabilities and the expression of each host gene. Genes were subsequently ranked according to their correlation coefficients, and the resulting ranked lists were used to perform Gene Set Enrichment Analysis (GSEA) of Gene Ontology (GO) Biological Process terms using the GSEApy Python library [29] through the Enrichr API [30]. Enriched terms were selected based on FDR *<* 0.05 and NES *>* 0, while over-representation analysis was additionally performed using unranked gene lists with an adjusted P-value threshold of 0.05.

### Datasets

PatHoST was evaluated using two high-resolution ST datasets representing complementary host–pathogen systems. These datasets were selected to assess the framework across different host species (mouse and human), pathogen classes (bacterium and virus), and experimental settings, providing a comprehensive benchmark for evaluating its ability to reconstruct pathogen niches from host gene expression.

#### Dataset 1 - Mouse model of *Mycobacterium abscessus* chronic infection

The primary benchmark dataset comprised nine tissue sections from murine model of chronic *Mycobacterium abscessus* lung infection, alongside control, generated using Visium Spatial Gene Expression platform. The dataset included seven infected lung sections and two sham-infected control. Among the infected tissues, four consecutive sections (*S*_1_*_−_*_4_) were processed with the dualST assay [2], two adjacent sections (*S_GT_* _1,2_) were profiled using the pathogen probe set only, and one section from an independent mouse was reserved as an external validation dataset (*S_EXT_* _1_). Four of these infected sections were generated for this study (*S*_3,4_ and *S_GT_* _1,2_), while the remaining two infected sections and the two sham-infected controls were retrieved from previously published data. The tissue sections underwent hematoxylin and eosin (H&E) staining using the Visium Spatial Gene Expression protocol designed for FFPE tissue deparaffinization, H&E Staining, Imaging and Decrosslinking demonstrated protocol. Imaging was performed using a scanning tile microscope (Aperio AT2, Leica). We exploited experimental procedures previously described [31] and animal studies were conducted according to protocols and adhering strictly to the Italian Ministry of Health guidelines for the use and care of experimental animals (IACUC N°1242) and approved by the San Raffaele Scientific Institute Institutional Animal Care and Use Committee (IACUC).

Additionally, two conventional ST sections from an independent *Klebsiella pneumoniae* infection model, profiled using host probes only, were included as specificity controls (*SK*1, 2) [32]. A comprehensive summary of all samples, experimental conditions, and data availability is provided in Supplementary Table S1. Detailed information regarding custom pathogen probe development, tissue processing, and preprocessing workflows is described in the following sections.

#### Experimental design for enhanced pathogen transcript detection

##### Custom pathogen probe design

Custom probes targeting *Mycobacterium abscessus* transcripts were designed following the 10x Genomics guidelines for Visium Spatial Gene Expression (Custom Probe Design for Visium Spatial Gene Expression and Chromium Single Cell Gene Expression Flex, CG000621). Candidate probe sequences were identified from the *M. abscessus* reference genome (ATCC 19977), targeting the *lsr2* and *rpoB* genes. Probe candidates were subsequently screened against reference genomes using BLAST [33] to ensure specificity for *M. abscessus* while excluding sequence similarity with *Homo sapiens*, *Mus musculus*, and other bacterial species. Three independent probe pairs were selected for each target gene according to the manufacturer’s design recommendations and details of the six custom pathogen probe are described by Di Marco & Nicola et al. [2].

##### Spatial transcriptomics data preprocessing

Raw Visium data were processed using the standard 10x Genomics analysis workflow with modifications to support custom pathogen probes. Tissue boundaries and fiducial alignment were manually refined using Loupe Browser v6.0.0, excluding empty spots corresponding to large airways. The resulting annotation files, together with the FASTQ files, were processed using Space Ranger v1.3.1 with default parameters. To enable the quantification of both host and *M. abscessus* transcripts, a custom reference was generated using the Space Ranger mkref utility by combining the mouse reference genome (mm10 Reference 2020-A) with the *M. abscessus* ATCC 19977 genome (NC 010397.1). The standard Visium Mouse Transcriptome Probe Set v1.0 was supplemented with the custom bacterial probes described above and used as the reference probe set during the spaceranger count workflow, following the 10x Genomics recommendations for custom probe design (CG000621).

The resulting Space Ranger outputs were imported into Scanpy [34] to generate separate host expression matrices and pathogen expression matrices, which served as the input data for all subsequent analyses performed by the PatHoST framework.

#### Dataset 2 - Human Lung tissues from SARS-CoV-2 infection

To evaluate the generalizability of PatHoST across host species and pathogen classes, we applied the framework to an independent ST dataset of human lung tissue infected with SARS-CoV-2, originally published by Sountart et al. [3]. Four infected lung sections from the COVID-19 cohort were analyzed together with six healthy human lung sections from Mayr et al. [17], which served as uninfected controls. Details of the selected samples are provided in Supplementary Table S4. The infected ST libraries targeted six SARS-CoV-2 genes, enabling the generation of host and viral expression matrices analogous to those used for the murine *M. abscessus* dataset.

### Statistical analysis

Statistical analyses were performed to compare methodological choices and evaluate the performance of alternative computational strategies. Comparisons involving continuous response variables were conducted using Student’s t-test or analysis of variance (ANOVA), as appropriate, whereas linear regression models were used to assess associations involving continuous predictors or combinations of continuous and categorical variables. When multiple statistical tests were performed, P-values were adjusted using the Bonferroni correction. Unless otherwise stated, statistical significance was defined as an adjusted *Pvalue <* 0.05.

## Code and Data availability

The PatHoST source code is publicly available at the GitHub repository: (Allen13x/PatHoST). The data can be requested to the authors.

## Author contributions

F.DM., A.G. and S.dP. conceived the study; A.G. and S.dP. supervised the project for the computational aspects; N.I.L. and S.dP. supervised the project for the experimental aspects; F.DM. lead the framework design, the implementation and the data analysis with the help of G.S.; F.N. lead the technical experimental activities; F.N. and N.I.L. evaluated the biological relevance of the findings; F.DM. S.dP. and A.G. wrote the manuscript with the help of F.N. and N.I.L. and the feedback from G.S. and D.M.C.; All authors approve the final manuscript.

## Acknowledgements

The authors thank A. Fiocchi and L. Guidotti (Animal Histopathology, IRCCS Ospedale San Raffaele, Milan, Italy) for the preparation of mouse histopathology samples, and the San Raffaele Microscopy Facility (Alembic, Ospedale San Raffaele, Milan, Italy) for histological processing and H&E image acquisition.

## Supplementary Materials

### Supplementary Tables

**Table S1:** Summary of murine lung section samples.

| Slice | Mouse ID | Infection | Lysozyme Condition | Host probes | Pathogen probes | Usage | Reference | GEO |
| --- | --- | --- | --- | --- | --- | --- | --- | --- |
| $S_1$ | 1 | *Mycobacterium abscessus* | A | Yes | Yes | Train | [2] | GSM7764368 |
| $S_2$ | 1 | *Mycobacterium abscessus* | A | Yes | Yes | Train | [2] | GSM8067258 |
| $S_3$ | 1 | *Mycobacterium abscessus* | A | Yes | Yes | Train | | |
| $S_4$ | 1 | *Mycobacterium abscessus* | C | Yes | Yes | Train | | |
| $S_{GT1}$ | 1 | *Mycobacterium abscessus* | B | No | Yes | Z-Projection | | |
| $S_{GT2}$ | 1 | *Mycobacterium abscessus* | D | No | Yes | Z-Projection | | |
| $S_{N1}$ | 2 | | A | Yes | Yes | Train | [2] | GSM7764369 |
| $S_{N2}$ | 2 | | A | Yes | Yes | Train | [2] | GSM8067259 |
| $S_{K1}$ | 3 | *Klebsiella pneumoniae* | / | Yes | No | Train | [32] | GSM5718440 |
| $S_{K2}$ | 3 | *Klebsiella pneumoniae* | / | Yes | No | Train | [32] | GSM5718442 |
| $S_{EXT1}$ | 1 | *Mycobacterium abscessus* | A | Yes | Yes | Test | | |

**Table S2:** Summary of the tested configurations for pathogen presence detection.

| Component | Tested Approaches |
| --- | --- |
| Preprocessing | Raw<br>Log normalization |
| Bacterial Beads regions | Kept<br>Discarded |
| Data Projection - Pathogen-only slice | No Donor<br>$S_{GT1}$<br>$S_{GT2}$<br>$S_{GT1} + S_{GT2}$ |
| Data Projection signal decay ($\lambda$) | [0.0, 0.5, 1] |
| Data Smoothing | No smoothing<br>Neighborhood Mean<br>Local PCA-Weighted Neighborhood Mean<br>Global PCA-Weighted Neighborhood Mean<br>MAGIC |
| Classifiers engines | Logistic Regression<br>Random Forest<br>XGBoost<br>Encoder<br>Autoencoder |
| Dropout rates - for encoders/decoders | [0.0 – 0.6] stepsize: 0.1 |

**Table S3:** Comparative performance analysis of PatHoST classifier engines. Area Under the Curve scores are reported for five model architectures across different validation stages. Performance was evaluated on the full training set, a subset restricted to infected slices, and an independent external dataset to test model generalizability. Values in bold indicate top-performing configurations.

| Model type | Training Dataset | Training Dataset - only Infected | External Dataset |
| --- | --- | --- | --- |
| Autoencoder | 0.85 | 0.64 | **0.86** |
| Encoder | 0.89 | 0.77 | **0.87** |
| Logistic Regression | 0.88 | 0.71 | **0.88** |
| XGBoost | **0.96** | **0.91** | 0.82 |
| Random Forest | **0.94** | **0.85** | **0.86** |

**Table S4:** Summary of human lung section samples.

| Slice | Infection | Usage | Reference | doi | ID |
| --- | --- | --- | --- | --- | --- |
| $C_1$ | *SARS-COV-2* | Train | [3] | 10.17632/xb2w7xvs2b.1 | 1135331 |
| $C_2$ | *SARS-COV-2* | Train | [3] | 10.17632/xb2w7xvs2b.1 | 1135332 |
| $C_3$ | *SARS-COV-2* | Train | [3] | 10.17632/xb2w7xvs2b.1 | 1112417 |
| $H_1$ | | Train | [17] | 10.5281/zenodo.10782711 | V19S23-092-A1 |
| $H_2$ | | Train | [17] | 10.5281/zenodo.10782711 | V10T03-282-A1 |
| $H_3$ | | Train | [17] | 10.5281/zenodo.10782711 | V10T31-015-A1 |
| $H_4$ | | Train | [17] | 10.5281/zenodo.10782711 | V10T31-019-A1 |
| $H_5$ | | Train | [17] | 10.5281/zenodo.10782711 | V10T03-280-A1 |
| $H_6$ | | Train | [17] | 10.5281/zenodo.10782711 | V10T03-281-A1 |
| $C_4$ | *SARS-COV-2* | Test | [3] | 10.17632/xb2w7xvs2b.1 | 1127396 |

**Table S5:** Comparative performance analysis of PatHoST classifier engines. Area Under the Curve scores are reported for five model architectures across different validation stages. Performance was evaluated on the full training set and an independent external dataset to test model generalizability. Values in bold indicate top-performing configurations.

| Model type | Training Dataset | Test slice |
| --- | --- | --- |
| Autoencoder | 0.94 | 0.61 |
| Encoder | 0.94 | 0.74 |
| Logistic Regression | **0.95** | 0.66 |
| XGBoost | 0.94 | 0.76 |
| Random Forest | **0.95** | **0.78** |

### Supplementary Figures

**Figure S1:**
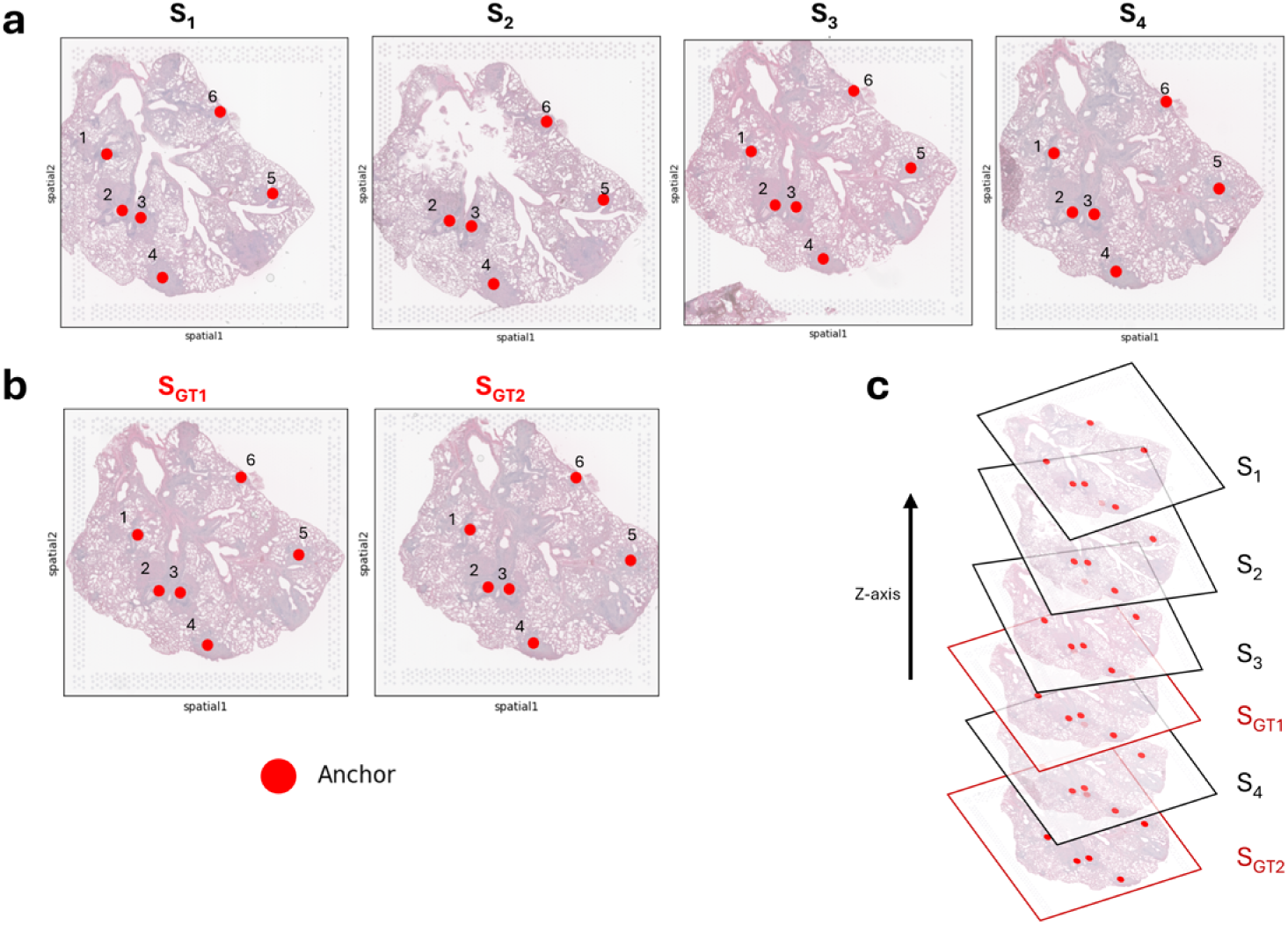
Characterization of Dataset #1 (murine lung chronically infected with *M. abscessus*), pathogen-only slices and z-alignment. (a) Slices *S*_1_*_−_*_4_ anchor points in red. (b) Slices *S_GT_* _1,2_ anchor points in red. (c) Z-stack of Slices *S*_1_*_−_*_4_ (black) and *S_GT_* _1,2_ (red). Anchor points in red.

**Figure S2:**
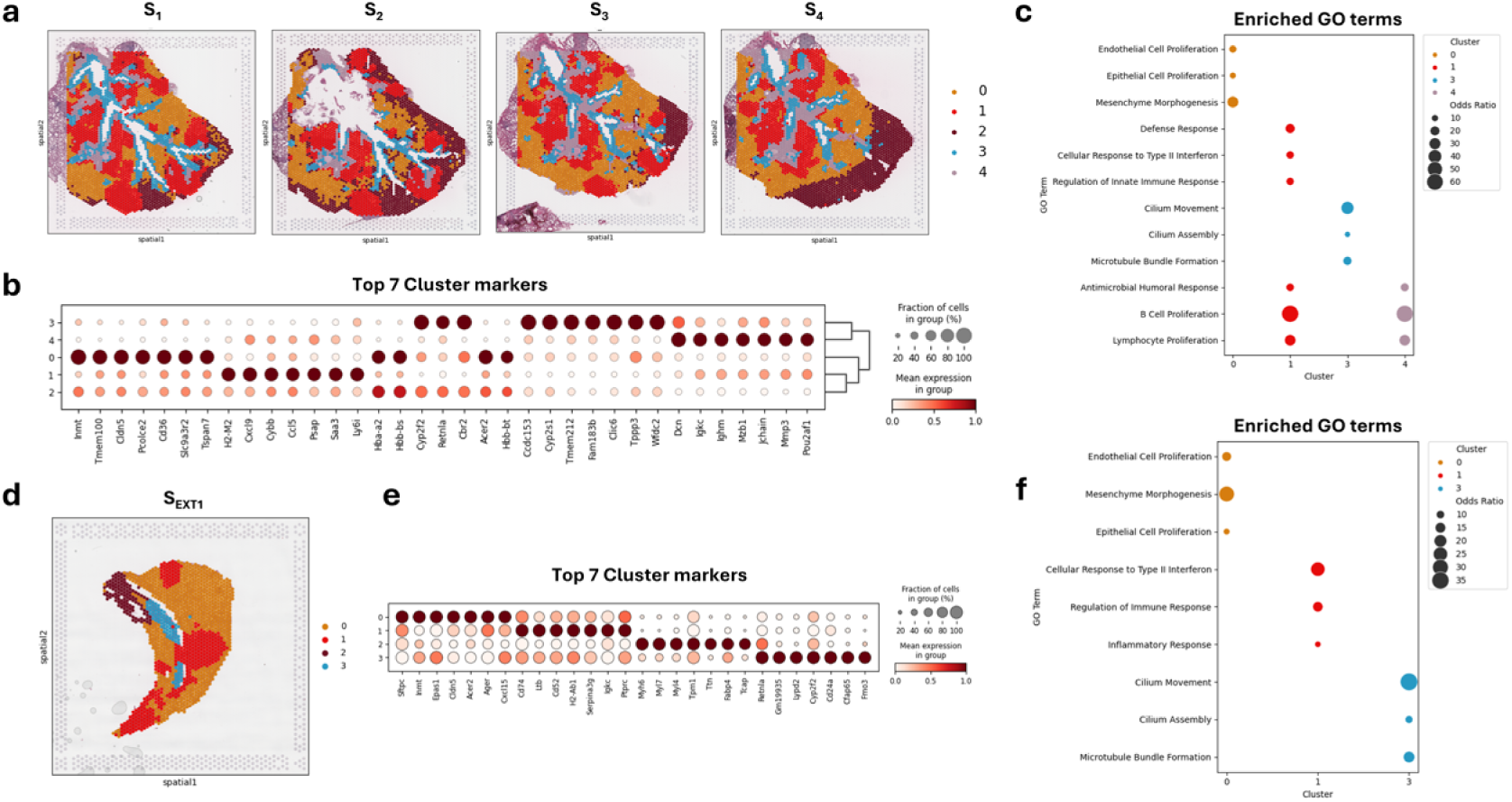
Characterization of Dataset #1 (murine lung chronically infected with *M. abscessus*), host transcriptome and clusters. (a) Slices *S*_1_*_−_*_4_ color-coded by host transcriptional clusters. (b) Dotplot showing the expression levels of the top seven transcriptional markers for each cluster of slices *S*_1_*_−_*_4_, as identified by differential expression analysis. The color scale represents the average log2 fold change when comparing the cluster to the others. The size represents the % of cells expressing the gene. (c) Dotplot displaying a selection of enriched Gene Ontology (GO) terms derived from cluster markers for slices *S*_1_*_−_*_4_. Dots are color-coded according to their respective clusters, and their size represents the odds ratio associated with each specific GO term (d) Slices *S_EXT_* _1_ color-coded by host transcriptional clusters. (e) Dotplot showing the expression levels of the top seven transcriptional markers for each cluster of slices *S_EXT_* _1_, as identified by differential expression analysis. The color scale represents the average log2 fold change when comparing the cluster to the others. The size represents the % of cells expressing the gene. Clusters markers are annotated according to biological functions on top. (f) Dotplot displaying a selection of enriched Gene Ontology (GO) terms derived from cluster markers for slices *S_EXT_* _1_. Dots are color-coded according to their respective clusters, and their size represents the odds ratio associated with each specific GO term.

**Figure S3:**
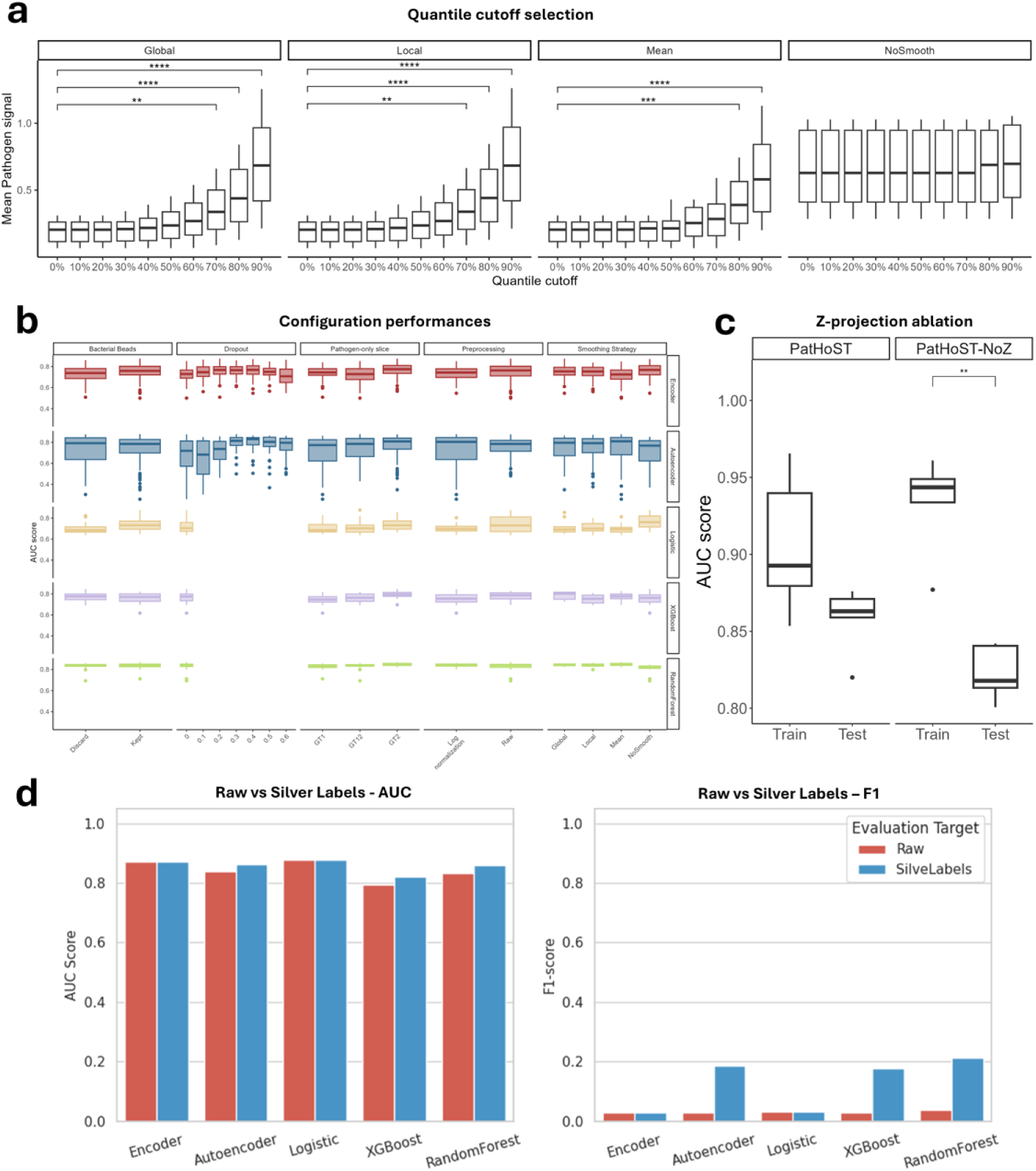
Characterization of Dataset #1 (murine lung chronically infected with *M. abscessus*) and PatHoST configurations performance. (a) Effect of quantile-based thresholding on Mean Pathogen signal distribution from infected slices (*S*_1_*_−_*_4_). Boxplots show the distribution of Mean Pathogen signal across different quantile cutoffs (0% to 90%) following z-projection. Results are faceted by smoothing method (Global, Local, Mean, and NoSmooth). Statistical comparisons indicate the divergence of filtered distributions relative to the baseline (0% cutoff); (b) Distribution of AUC scores across various model configurations on external slice (*S_EXT_* _1_). Data are faceted by preprocessing parameters and smoothing strategies. Different colors and rows represent the specific models evaluated; (c) Boxplots showing AUC performance on the external dataset (*S_EXT1_*) for PatHoST (Z-projection + Smoothing) and the PatHoST-NoZ (only Smoothing module) approach across all training data. (d) Barplots for comparison of model classification performance evaluated against highly sparse pathogen Raw Counts (red) and spatially imputed Silver Labels (blue) on the external validation slice (*S_EXT1_*). On the left, the AUC score shows consistently high values across raw and silver labels. On the right, F1 scores show inconsistency between raw and silver labels, and low values due to the presence of unlabeled positive spots due to technical dropouts.

**Figure S4:**
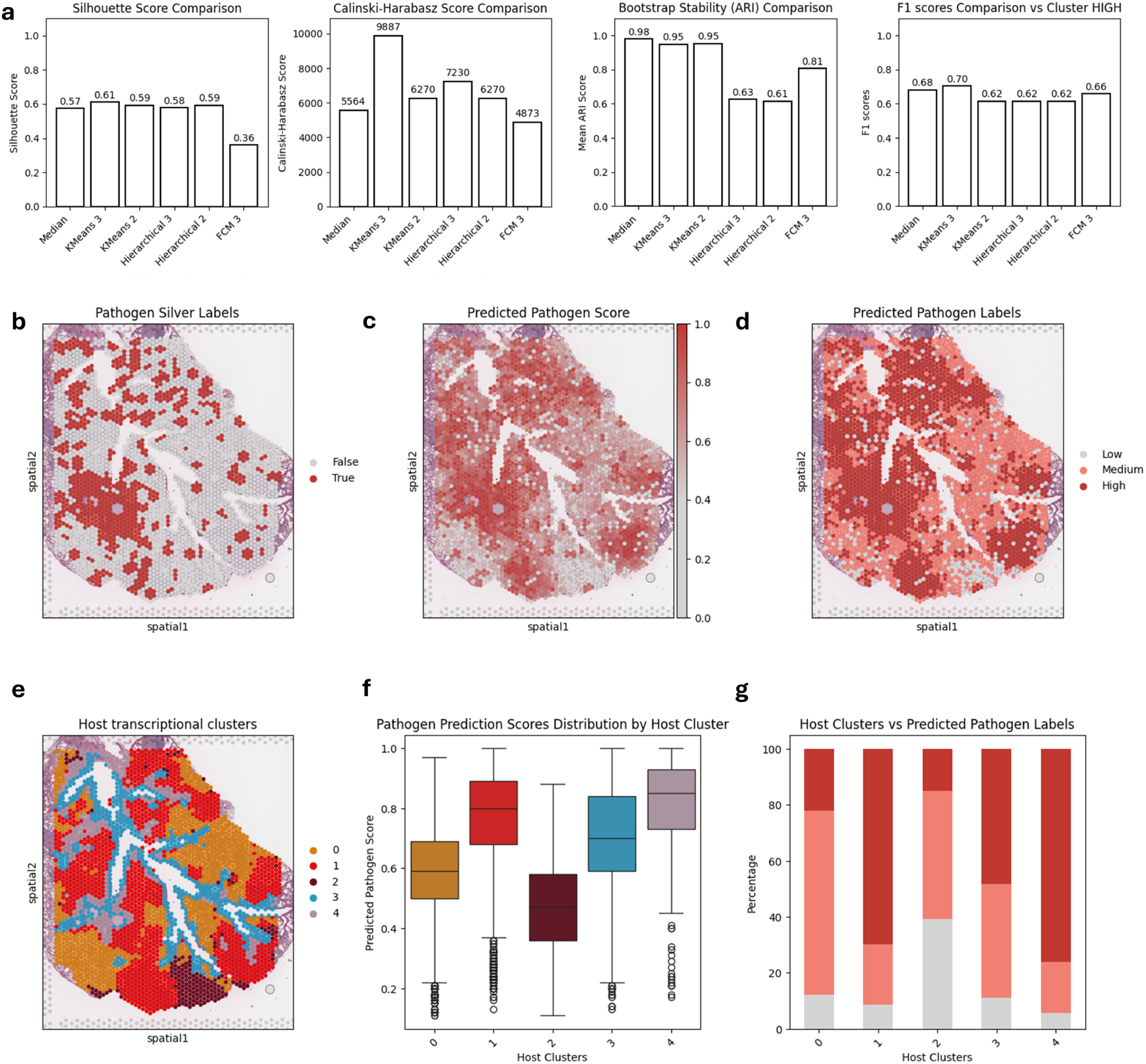
Evaluation of discretization strategies on Dataset #1 infected slices (*S*_1_*_−_*_4_) (a) Values for Silhouette, Calinski-Harabasz, Adjusted Rand Index, and F1 scores for Pathogen labels obtained with different thresholding methods from PatHoST predicted pathogen scores on infected slices (*S*_1_*_−_*_4_); (b) Pathogen silver label on slice *S*_1_ after Z-projection and Global neighborhood PCA based smoothing; (c) Predicted Pathogen scores on slice *S*_1_ produced by a Random Forest model; (d) Predicted Pathogen labels on slice *S*_1_ using k-mer algorithm with *k* = 3; (e) Spatial map of host transcriptional clusters identified in *S*_1_; (f) Distribution of predicted pathogen scores across transcriptional clusters; (g) Distribution of predicted pathogen labels across transcriptional clusters.

